# Identification and Characterization of Long Non-Coding RNA in Tomato Roots under Salt Stress

**DOI:** 10.1101/2021.11.12.468377

**Authors:** Ning Li, Zhongyu Wang, Baike Wang, Juan Wang, Ruiqiang Xu, Tao Yang, Shaoyong Huang, Qinghui Yu, Huan Wang, Jie Gao

## Abstract

As one of the most important vegetable crops in the world, the production of tomatoes was restricted by salt stress. Therefore, it is of great interest to analyze the salt stress tolerance genes. As the non-coding RNAs (ncRNAs) with a length of more than 200 nucleotides, long non-coding RNAs (lncRNAs) lack the ability of protein-coding, but they can play crucial roles in plant development and response to abiotic stresses by regulating gene expression. Nevertheless, there are few studies on the roles of salt-induced lncRNAs in tomatoes. Therefore, we selected wild tomato *Solanum pennellii* (*S. pennellii*) and cultivated tomato M82 to be materials. By high-throughput sequencing, 1044 putative lncRNAs were identified here. Among them, 154 and 137 lncRNAs were differentially expressed in M82 and *S. pennellii*, respectively. Through functional analysis of target genes of differentially expressed lncRNAs (DE-lncRNAs), some genes were found to respond positively to salt stress by participating in Abscisic Acid (ABA) signaling pathway, brassinosteroid (BR) signaling pathway, ethylene (ETH) signaling pathway and anti-oxidation process. We also construct a salt-induced lncRNA-mRNA co-expression network to dissect the putative mechanisms of high salt tolerance in *S. pennellii*. We analyze the function of salt-induced lncRNAs in tomato roots at the genome-wide levels for the first time. These results will contribute to understanding the molecular mechanisms of salt tolerance in tomatoes from the perspective of lncRNAs.

## 1. Introduction

As the second-largest vegetable in the world, there is a growing demand for tomatoes. However, the area of the world’s saline-alkali land has increased annually, soil salinization has become a major impediment to affect the growth and development of tomatoes and reduce tomato production seriously [1]. Previous studies have demonstrated that salt stress usually causes ion stress [2], osmotic stress [3] and secondary damage to plants [4], especially oxidative stress damage [5, 6]. Therefore, in order to adapt to salt stress, plants need to rebuild the homeostasis of cell ions, osmosis, and redox balance [7]. Researches on important signaling pathways for salt tolerance have been the subject of many classic studies. So far, scientists have revealed many important signaling pathways of salt tolerance in plants. Such as the Salt Overly Sensitive (SOS) signal pathway, mitogen-activated protein kinase (MAPK) cascade signal pathway, CDPK cascade reaction pathway, ABA signal pathway and so forth. These results reveal that the molecular mechanisms resulting from salt stress tolerance are very complex in plants. Therefore a thorough understanding of the salt tolerance mechanism is crucial.

In recent years, the role of non-coding RNA (ncRNA) in plants has attracted more and more attention. Quite a few studies have shown that ncRNA plays a crucial role in different biological processes of plants, such as cell development [8], regulation of epigenetics [9], transcription and translation [10] and so forth. Numerous studies show that ncRNAs have critical roles in diverse biological processes from plants to animals, such as sponging by microRNAs, cell development, acting as modular scaffolds, and regulating epigenetic inheritance. The ncRNAs include rRNA, tRNA, snRNA, snoRNA, miRNA, lncRNA, and other RNAs with known functions. Long non-coding RNAs (lncRNAs) is a large class of transcripts from the non-protein coding region of the genome that contain more than 200 nucleotides but lack protein coding ability, and do not contain or contain short open reading frame (ORF), usually exert a regulatory role in the response of plants to abiotic stress [11]. According to the genomic positioning of its coding genes to adjacent protein-coding genes, lncRNAs can be further divided into long intergenic non-coding RNAs (lincRNAs), natural antisense transcripts (NATs), and intronic RNAs (incRNAs). With the development of technology, many lncRNAs transcripts have been identified in different plant species by tiling array and transcriptome reassembly, like *Arabidopsis* [12], rice [13, 14], soybean [15, 16] and cotton [17, 18] and so forth. Considering the complexity of lncRNA regulation, there are merely a few functional characteristics of lncRNAs in plants so far, but in recent years, researches on the specific role of lncRNA in plants have become more and more attractive.

Recent evidence suggests that lncRNAs play essential roles in tomatoes during flowering [19], resistance to *Phytophthora infestans* [20, 21], fruit ripening [22], resistance to drought [23], and are also studied in *Arabidopsis thaliana* [24], *Camellia sinensis* [25], *Gossypium hirsutum* [26]. In recent years, competitive endogenous RNA (ceRNA) has provided an innovative way to study the molecular mechanisms of stress in plants. Among the drought stress-related lncRNAs of *Populus tomentosa*, some lncRNAs are identified as competitive endogenous RNAs, which can combine with known poplar miRNAs to regulate the expression of miRNAs target genes. At the same time, the qRT-PCR result had also verified this result [27]. Since miR398 can respond to different stresses, tae-miR398 regulates low-temperature tolerance by down-regulating its target gene CSD1 in the cold hardiness mechanism of winter wheat. LncRNA can indirectly regulate the expression of CSD1 by competitively binding miR398, thereby affect the cold resistance of Dn1. The regulation of miR398 triggers a regulatory loop that is essential for cold resistance in wheat [28].

The role of lncRNAs in the process of tomato salt stress is rarely studied. However, what is not yet clear is the importance of salt-induced lncRNA in tomatoes. In comparison with cultivated tomato, wild tomato display increased salt tolerance and has stronger salt tolerance. Due to different growing environments and reproductive isolation, some salt-responsive genes in wild tomatoes may not exist in cultivated tomato species. To use modern molecular biology techniques to improve the salt tolerance of cultivated tomatoes, it is first necessary to understand the molecular mechanism of tomato salt tolerance, and wild tomatoes are important germplasm resources for revealing the salt tolerance mechanism and mining salt tolerance genes. Consequently, in this study, we selected wild tomato *Solanum pennellii* (*S. pennellii*) and cultivated tomato M82 (*Solanum Lycopersicum L.*) as materials. The functions of salt-induced lncRNAs in the two cultivars are analyzed and compared by analyzing the target genes. This essay attempts to show that some salt-related genes might be regulated by salt-induced lncRNAs in *S. pennellii.* We also construct the corresponding salt-induced co-expression network. In general, this paper presents new evidence for the putative mechanism of salt tolerance in tomatoes from the perspective of the lncRNA-mRNA network.

## 2. Results

### 2.1. Genome-Wide Identification and Characterization of LncRNAs in Tomatoes

To identify the lncRNAs in cultivated tomato and wild tomato in response to salt stress, systematically. In this study, we used roots of wild tomato and cultivated tomato under salt stress for 12 hours to be materials, and then performed the whole transcriptome sequencing analysis. Each sample has three replicates. The prepared library was sequenced on the Illumina-Hiseq platform, and finally a total of 588.98G reads were obtained. By removing reads containing adapter, reads containing ploy-N and low-quality reads from raw data, clean reads were eventually obtained (Supplementary Table S1). According to the analysis procedure in Figure 1A, we identified 1044 putative lncRNAs (Supplementary Table S2), which were distributed on all chromosomes of tomato. Among them, the number of lncRNAs on chromosome 1 had the most lncRNAs (117 lncRNAs) and the number of lncRNAs on chromosomes 2 and 6 were the least (only 58 lncRNAs) (Figure 1B). As shown in Figure 1B, the density distribution of lncRNAs predicted in tomatoes was almost uniform with little difference. LncRNAs had the highest density on chromosome 7 (~1.5 lncRNAs/Mbp nucleotides) and the lowest density on chromosome 3 (~0.91lncRNAs/Mbp nucleotides). The 1044 lncRNAs include 859 lincRNAs, 165 antisense lncRNAs, 11 exonic, and 9 intronic lncRNAs (Figure 1C). Through further analysis of the length of these lncRNAs, it was found that the majority of them were shorter than 2000 nt (Figure 1D) and the mean length of lncRNAs was shorter than that of mRNAs as a whole.

**Figure 1.**
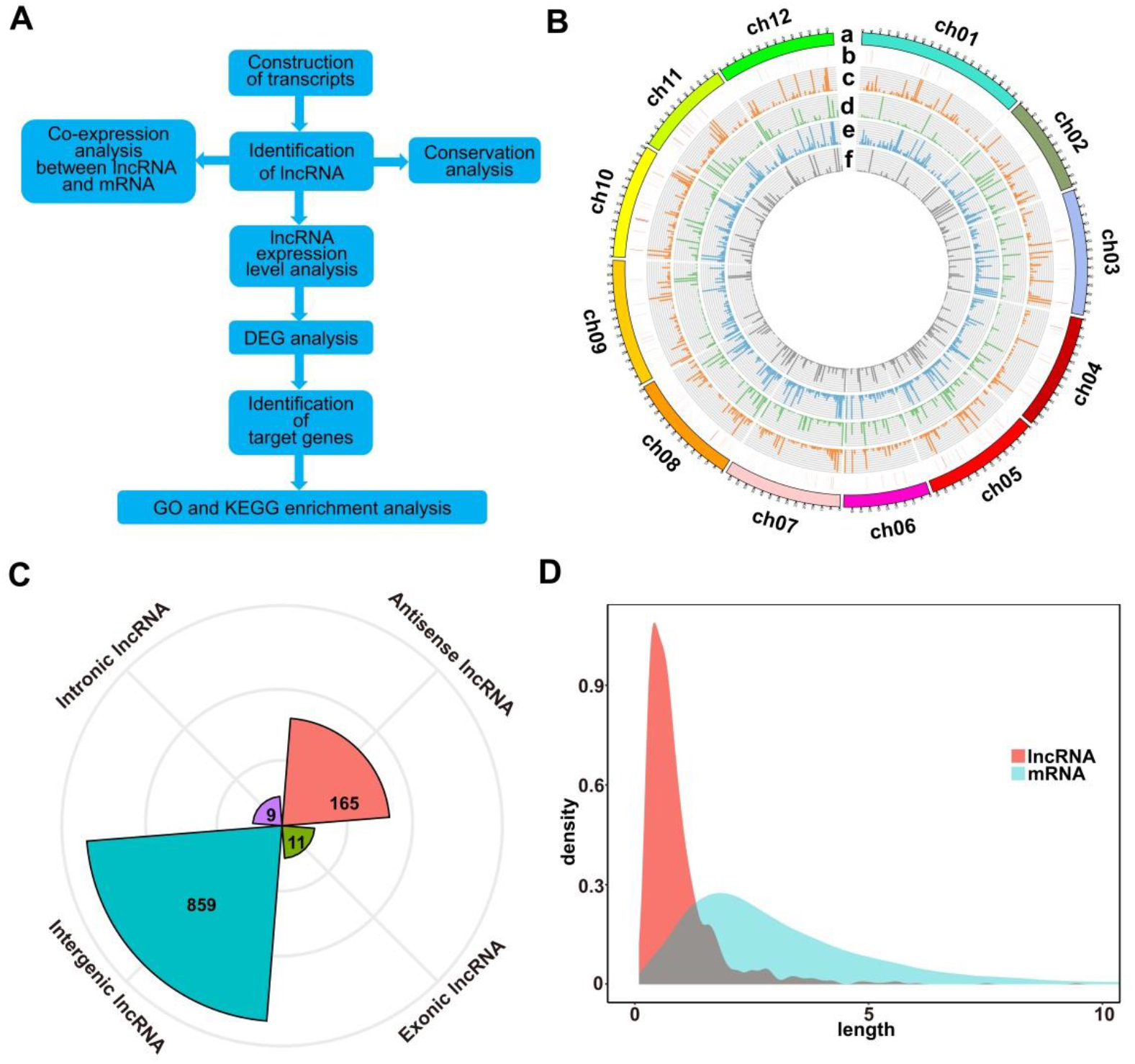
Identification and Characterization of lncRNAs in tomatoes. (**A**) The pipeline for the identification of lncRNAs in tomatoes. (**B**) Circos plot depicting the distribution and expression of identified lncRNAs. From outer to inner circles, a-f circles represent chromosomes, lncRNAs distribution on chromosomes, the expression levels of lncRNAs in the control sample of M82, the expression levels of lncRNAs in the control sample of *S. pennellii*, the expression levels of lncRNAs in the salt-treated sample of M82, the expression levels of lncRNAs in the salt-treated sample of *S. pennellii*, respectively. The vertical lines represent the high-low values of lncRNAs (**C**) Radar chart represents the numbers of the four lncRNA types. (**D**) Length distribution of lncRNAs and mRNAs, orange represents lncRNAs, blue represents mRNAs.

### 2.2. Identification of Salt-Responsive DE-lncRNAs

To compare and analyze the different lncRNAs in response to salt stress in cultivated tomato and wild tomato, we compared and analyzed the expression levels of lncRNAs in response to salt stress in the two cultivars. There were 406 and 8 lncRNAs expressed specifically in M82 and *S. pennellii*, respectively. 630 lncRNAs showed expression in both cultivars (Figure 2A, Table S2). And the average expression levels of lncRNAs were lower than that of mRNAs (Figure 2B). Subsequently, comparing the standardized expression of mRNAs between cultivated and wild tomatoes, the significant differences were also founded by clustering (Figure 2C). In M82, 154 lncRNAs were significantly differential expressed, of which 133 were up-regulated and 21 were down-regulated (Supplementary Table S3). In contrast, 137 lncRNAs were differentially expressed in *S. pennellii*, of which 37 were up-regulated and 100 were down-regulated (Figure 2C, Supplementary Table S4). It could be seen that the expression levels of most DE-lncRNAs were significantly down-regulated in *S. pennellii*, while the expression levels of most differentially expressed lncRNAs (DE-lncRNAs) in M82 were significantly up-regulated. Only 33 lncRNAs were differentially expressed both in the two cultivars, of which 15 lncRNAs were expressed at opposite levels (Fig2C, D). Interestingly, the expression levels of these 15 lncRNAs were all up-regulated in M82, while their expression levels were down-regulated in *S. pennellii*, indicating that these 15 lncRNAs might be relevant to the high salt tolerance of *S. pennellii*.

**Figure 2.**
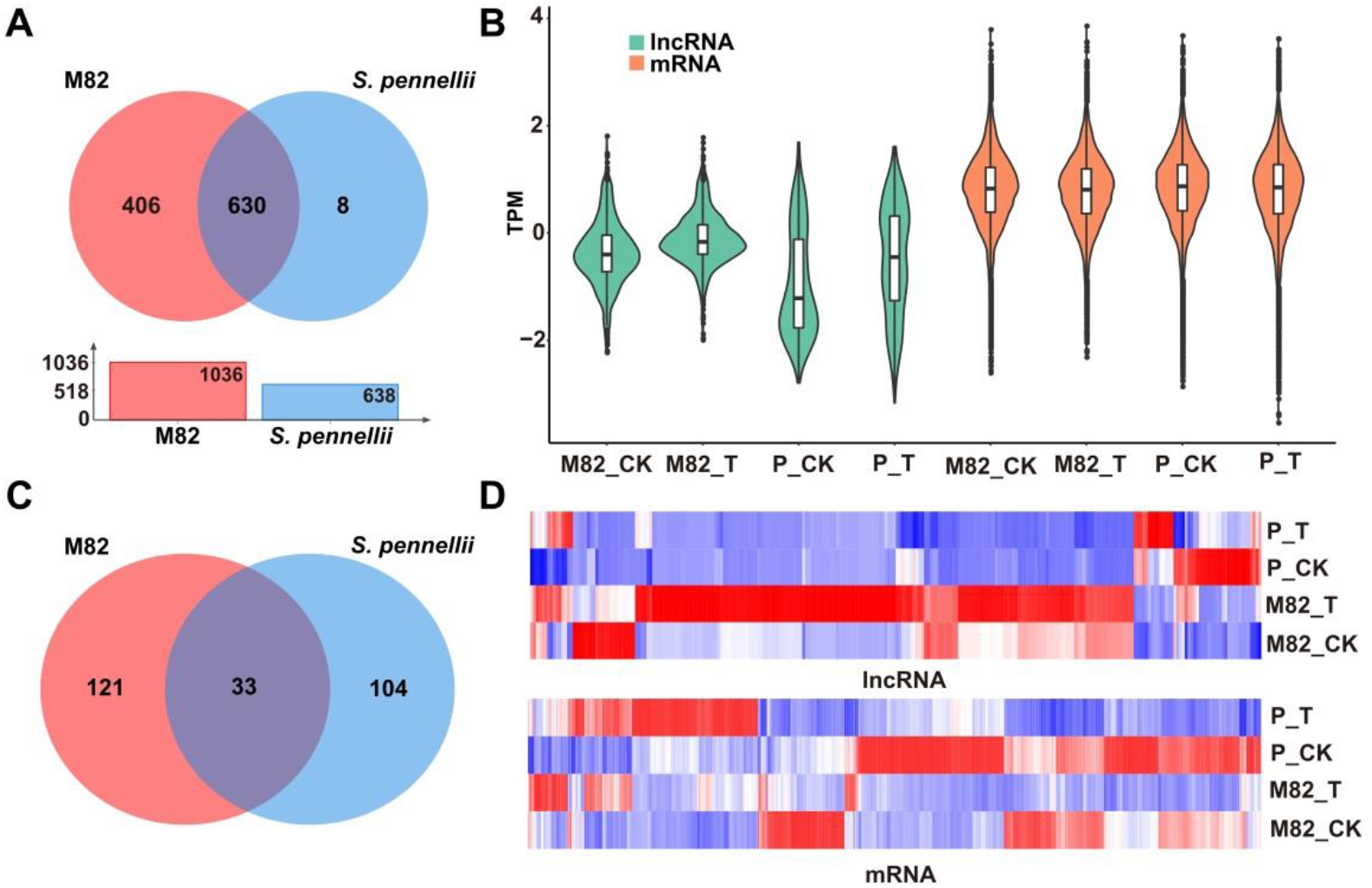
Expression patterns analysis of lncRNAs and mRNAs. (**A**) Venn diagram of common and specific lncRNAs. Red represents M82, blue represents *S. pennellii.* The overlap represents the lncRNAs that were expressed in both M82 and *S. pennellii*. The red bar and blue bar in the bar graph represent the number of lncRNAs that were expressed in M82 and *S. pennellii*, respectively. (**B**) Violin plot of expression levels for lncRNA and mRNA transcripts (showed in the relative expression level of mRNA/lncRNA presented by log-transformed). (**C**) The Venn diagram shows significantly DE-lncRNAs (P-value <0.05). Overlaps represent the lncRNAs that were differentially expressed in both M82 and *S. pennellii*. (**D**) Heatmap presentation of relative expression levels of differentially expressed lncRNAs and mRNAs. Z-score represents the regulation trends, red represents up-regulation, blue represents down-regulation. The color scale bar shows z-score values after z-score row normalization.

### 2.3. Cis-Regulation of LncRNA Neighboring Genes

To further obtain the possible biological function of lncRNAs in tomatoes under salt stress, we treated the genes within 1 Mbps upstream/downstream DE-lncRNAs as cis-regulated genes and performed GO and KEGG analysis based on these genes. The results revealed that 125 DE-lncRNAs targeted 1227 differential expressed mRNAs (DE-mRNAs) in M82 and 111 DE-lncRNAs in *S. pennellii* target 1268 DE-mRNAs. In the end, we obtained 1612 and 1535 pairs of lncRNA-mRNA genes in M82 and *S. pennellii*, respectively. Among them, 700 pairs in M82 were positively correlated with expression levels, and 912 pairs were negatively correlated with expression levels (Supplementary Table S5). In *S. pennellii*, 880 pairs of expression levels are positively correlated, and 655 pairs of expression levels are negatively correlated (Supplementary Table S6).

In M82, the GO analysis results of these co-localized genes showed that 13, 13, 9 and 10 genes were significantly enriched in DNA replication (GO:0006260), cell wall macromolecule metabolic process (GO:0044036), xyloglucan metabolic process (GO:0010411), hemicellulose metabolic process (GO:0010410) and other terms (Figure 3A, Supplementary Table S7). In *S. pennellii*, the GO terms that were significantly enriched contained cell wall macromolecule metabolic process (GO:0044036), oxidation-reduction process (GO:0055114), cell wall organization or biogenesis (GO:0071554), and photosynthesis, light harvesting (GO:0009765) (Figure 3B, Supplementary Table S8). Compared with M82, there were more genes enriched in the oxidation-reduction process in *S. pennellii*, which might indicate that the steady-state of redox balance in *S. pennellii* is higher than that in M82 under salt stress.

**Figure 3.**
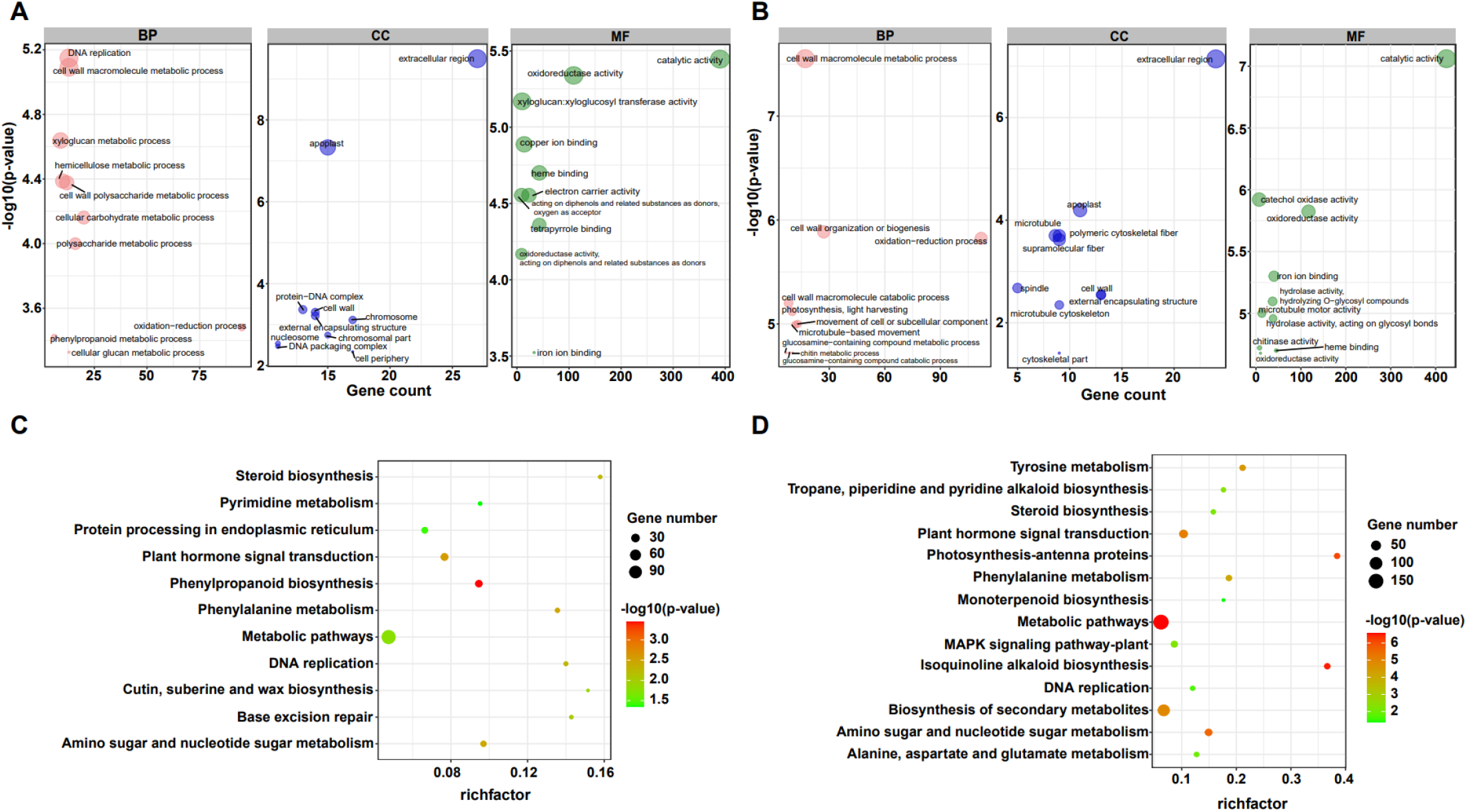
GO and KEGG enrichment analysis of the cis-regulated target genes of lncRNAs. (**A**) The top 10 significant terms in Biological Process (BP), Molecular Function (MF), and Cellular Component (CC) of GO enrichment analysis (P-value < 0.05) in M82. The larger the circle, the more significant of enrichment of the target genes in this pathway is. (**B**). The top 10 significant terms in Biological Process (BP), Molecular Function (MF), and Cellular Component (CC) of GO enrichment analysis (P-value < 0.05) in *S. pennellii*. (**C**) The KEGG enrichment scatters plot of cis-regulated target genes of lncRNAs in M82. The size of the circles represents the number of genes, the color of the circle represents the P-value. The abscissa represents the rich factor, the larger the rich factor, the greater the degree of enrichment (RichFactor is the ratio of the number of differentially expressed genes to the number of all genes in this pathway term). (**D**) The KEGG enrichment scatters plot of cis-regulated target genes of lncRNAs in *S. pennellii*.

KEGG analysis results showed that these co-localized genes in M82 and *S. pennellii* were significantly enriched into 11 and 14 KEGG pathways, respectively. among them, Phenylpropanoid biosynthesis, Base excision repair, Cutin, suberine and wax biosynthesis, Protein Processing in endoplasmic reticulum and Pyrimidine metabolism were only enriched in M82 (Supplementary Table S9), Isoquinoline alkaloid biosynthesis, Photosynthesis-antenna proteins, Biosynthesis of secondary metabolites, Tyrosine metabolism, Tropane, piperidine and pyridine alkaloid biosynthesis, MAPK signaling pathway-plant, Alanine, aspartate and signaling pathway metabolism and Monoterpenoid biosynthesis were only enriched in *S. pennellii* (Supplementary Table S10). Plant hormone signal transduction, Phenylalanine metabolism, Amino sugar and nucleotide sugar metabolism, DNA replication and Steroid biosynthesis were enriched in *S. pennellii*. The number of genes enriched in these pathways was more than that of M82.

### 2.4 Trans-Regulation of Target Genes by LncRNAs

LncRNAs could affect the expression of distant genes by regulating remote mRNAs transcription. Therefore, in order to identify trans-regulated genes of DE-lncRNA that might be involved in the response to salt stress, we analyzed the correlation between DE-lncRNAs and DE-mRNAs (correlation coefficient r ≥ 0.99 or ≤ −0.99). In M82, we predicted that 3106 genes had a trans-regulatory relationship with 145 lncRNAs, of which 14922 pairs are positively correlated, and 7341 pairs are negatively correlated (Supplementary Table S11). In *S. pennellii*, it was predicted that 3244 genes have a trans-regulatory relationship with 118 lncRNAs, of which 14126 pairs were positively correlated, and 6261 pairs were negatively correlated (Supplementary Table S12). These lncRNA-mRNA gene pairs were all significantly correlated (p<0.05).

By comparing the expression levels of these target genes, we found that 1685 and 1823 genes were regulated by lncRNAs only in M82 and *S. pennellii*, respectively. There were 1421 genes regulated by lncRNAs in both two cultivars. Among them, the expression levels of 47 genes exhibited the opposite trend, including 24 up-regulated genes in *S. pennellii* and 23 down-regulated genes in M82. For example, Solyc06g074710.1 (Hydroxyacid Hydroxycinnamoyltransferase, HCT), Solyc04g005810.3 (Thioredoxin-h2, TRXH2), Solyc08g065500.2 (Protein phosphatase 2C, PP2C), Solyc08g065670.3 (PP2C), Solyc11g010400.2 (2-oxoglutarate, 2-OG) and Solyc11g010400.2 (Fe (II)-dependent oxygenase, DMR6) were significantly up-regulated in *S. pennellii* and significantly down-regulated in M82.

In M82, Solyc04g005810.3 (TRXH2) could be targeted by 9 lncRNAs, and their expression patterns showed a negative correlation. Two PP2C genes (Solyc08g065500.2 and solyc08g065670.3) could be targeted by lnc_000557 at the same time and exhibited the same trend in expression. Solyc11g010400.2 (DMR6) could be targeted by 4 lncRNAs (Lnc_000288, Lnc_000758, Lnc_000964 and Lnc_001011), of these, Lnc_000288 could also target Solyc06g074710.1 (HCT). In addition to the Lnc_001011-Solyc11g010400.2 pair, the other four pairs showed the same trend in expression. In *S. pennellii*, Solyc06g074710.1 (HCT) and Solyc11g010400.2 (DMR6) could be targeted by 9 lncRNAs simultaneously, and the same trend was seen between these two genes and 9 lncRNAs were the same. Beyond this, these two genes could also be targeted by two different lncRNAs (Lnc_000364-Solyc06g074710.1, Lnc_000837-Solyc06g074710.1, Lnc_000839-Solyc11g010400.2, Lnc_000996-Solyc11g010400.2). With the exception of the Lnc_000837-Solyc06g074710.1 pair, the other 3 pairs showed the opposite trend in expression. In common with Solyc06g074710.1 (HCT), Solyc04g005810.3 (TRXH2) could be targeted by 3 lncRNAs (Lnc_000334, Lnc_000364 and Lnc_000600). Except for Lnc_000364-Solyc04g005810.3 (negative correlation), the expression levels of the other two pairs showed remarkable positive correlations. In contrast to M82, two PP2C genes (Solyc08g065500.2 and solyc08g065670.3) could be targeted by 3 lncRNAs (Lnc_000181, Lnc_000359, and Lnc_000764) and showed remarkable positive correlations in *S. pennellii*.

To further investigate the potential function of lncRNAs, we divided the target genes of lncRNAs into 3 sets: the target genes that were regulated by lncRNAs only in M82 and *S. Pennellii*, respectively, the target genes that were regulated by lncRNAs in both two cultivars. By performing the GO and KEGG enrichment analysis of these 3 sets of target genes, the differences of salt-responsive lncRNAs in two cultivars were compared. The GO enrichment results were as follows. There were 1685 genes targeted by lncRNAs only in M82, among them, 285 genes (112 up-regulated genes and 173 down-regulated genes) were significantly enriched in 10 GO terms (Supplementary Table S13), including oxidation-reduction process (GO: 0055114), hydrogen peroxide metabolic process (GO: 0042743), reactive oxygen species metabolic process (GO: 0072593), response to oxidative stress (GO: 0006979), Transport (GO: 0006810), Response to Chemical (GO: 0042221), and so forth. There were 1823 genes targeted by lncRNAs only in *S. pennellii*, among them, 381 genes (108 up-regulated genes and 273 down-regulated genes) were significantly enriched in 10 GO terms (Supplementary Table S14), including photosynthesis, light harvesting (GO:0009765), photosynthesis (GO:0015979), movement of cell or subcellular component (GO:0006928), microtubule-based movement (GO:0007018), microtubule-based process (GO:0007017), carbohydrate metabolic process (GO:0005975), oxidation-reduction process (GO:0055114), generation of precursor metabolites and energy (GO:0006091), response to stimulus (GO:0050896), hydrogen peroxide metabolic process (GO:0042743). Interestingly, there were 3 GO terms associated with photosynthesis, which contained 37 differentially expressed genes. Except for Solyc12g017250.2 (photosystem II subunit R, PSBR), which was up-regulated, the other 36 genes were significantly down-regulated in *S. pennellii*. Moreover, no significant change in expression levels of the 37 photosynthesis-related genes in M82, the expression levels of these genes in M82 were also lower than that of these genes in *S. pennellii*, significantly. The 37 photosynthesis-related genes could be targeted by 37 lncRNAs. There were 1421 genes targeted by lncRNAs in both two cultivars. For these genes, cell wall biogenesis metabolic process (GO:0042546 and GO:0044036), cell wall polysaccharide metabolic process (GO:0010383), oxidation-reduction process (GO:0055114), glucan metabolic process (GO:0044042), xyloglucan metabolic process (GO:0010411) were enriched (Figure S1, Supplementary Table S15).

In addition, the GO term oxidation-reduction process contained 142 and 148 genes in M82 and *S. pennellii*, respectively. In M82, 58 up-regulated and 84 down-regulated genes were significantly enriched, and 49 up-regulated and 99 down-regulated genes were enriched in *S. pennellii*. The GO term response to stimulus contained 97 and 118 genes in M82 and *S. pennellii*, respectively. Interestingly, the 97 genes in M82 were only differentially expressed in M82 and shown no significant change in *S. pennellii*. However, there were 40 genes only up-regulated differentially in *S. pennellii* among the 118 genes, which consisted of Solyc09g090970.3 (MLP-like protein 423, MLP423), Solyc12g056650.2 (GIGANTEA, GI), Solyc05g052270.2 (calcium-independent protein kinase 10, CIPK10), Solyc12g010130.1 (CIPK6), Solyc08g068960.3 (histidine kinase 5, HK5), Solyc02g083620.3 (ascorbate peroxidase 3, APX3), Solyc08g067310.1 (CIPK5) and some efflux protein genes. In *S. pennellii*, these 40 genes could be targeted by 27 lncRNAs with negative regulatory relationships between 10 lncRNAs and their target genes and the other 17 lncRNAs showed positive regulatory relationships with their target genes.

KEGG pathway analysis revealed that there were 197 genes (82 up-regulated genes and 115 down-regulated genes) were enriched in Nitrogen metabolism, Phenylpropanoid biosynthesis, Plant-pathogen interaction, Biosynthesis of secondary metabolites, Amino sugar and nucleotide sugar metabolism, Plant hormone signal transduction, and several metabolic pathways in M82 (Supplementary Table S16). In particular, there were 17 genes enriched in the Plant hormone signal transduction pathway up-regulated significantly. In *S. pennellii*, 344 genes (105 up-regulated genes and 239 down-regulated genes) were significantly enriched in Photosynthesis-antenna proteins, Biosynthesis of secondary metabolites, Phenylpropanoid biosynthesis, Photosynthesis, Amino sugar and nucleotide sugar metabolism, Plant hormone signal transduction, Carbon fixation in photosynthetic organisms, MAPK signaling pathway-plant and so on (Supplementary Table S17). There were 57 genes significantly enriched in Photosynthesis-related pathways and most of them were down-regulated in *S. pennellii*. There were also 12 genes enriched in Plant hormone signal transduction and significantly down-regulated. Known from the comparison of KEGG results, Nitrogen metabolism, Plant-pathogen interaction, Base excision repair, Pyrimidine metabolism, and beta-Alanine metabolism were enriched only in M82 and Photosynthesis-antenna proteins, Photosynthesis, Carbon fixation in photosynthetic organisms, and MAPK signaling pathway-plant were enriched only in *S. pennellii* (Fig4C, D). In M82, 13 genes were enriched in the Nitrogen metabolism term, of which 11 genes were down-regulated. It showed that the nitrogen metabolism pathway was severely affected. In *S. pennellii*, 22 genes were enriched in MAPK signaling pathway-plant, of which 12 genes were up-regulated significantly. 143 genes were enriched in the Biosynthesis of secondary metabolites, of which 51 genes were up-regulated significantly. In addition, Phenylalanine metabolism, MAPK signaling pathway, Plant hormone signal transduction, Phenylpropanoid biosynthesis, and Biosynthesis of secondary metabolites pathways were enriched in both two cultivars (Figure S2, Supplementary Table S18). These results illustrated that the salt-responsive pathways in M82 and *S. pennellii* were significantly different, lncRNA might be involved in the response to salt stress through regulating their potential target genes. Apart from that, these lncRNAs might play an essential role in enhancing salt tolerance in *S. pennellii*.

**Figure 4.**
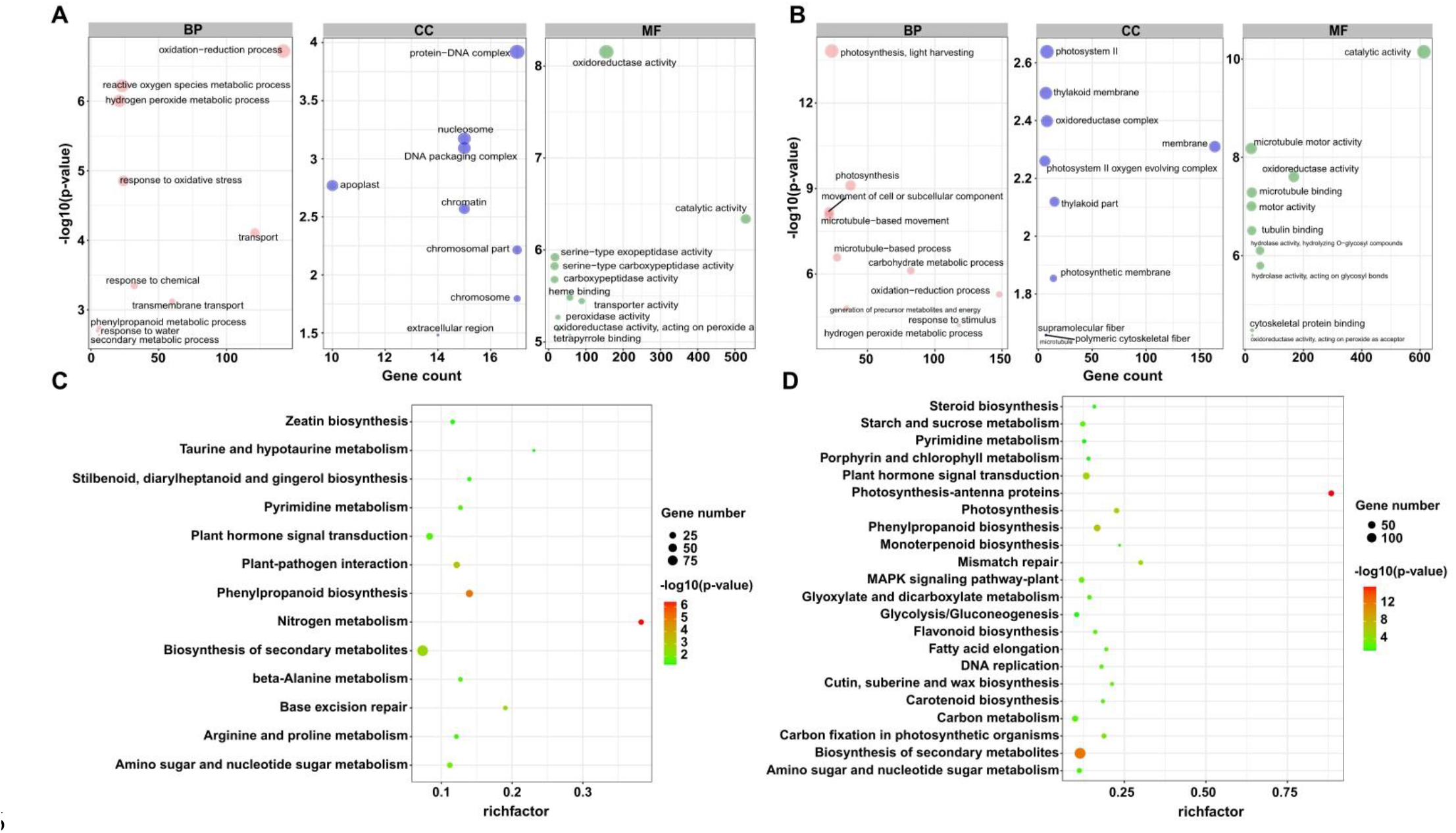
GO and KEGG enrichment analysis of the trans-regulated target genes of lncRNAs. (**A**) The top 10 significant terms in Biological Process (BP), Molecular Function (MF), and Cellular Component (CC) of GO enrichment analysis (P-value < 0.05) in M82. The larger the circle, the more significant of enrichment of the target genes in this pathway is. (**B**). The top 10 significant terms in Biological Process (BP), Molecular Function (MF), and Cellular Component (CC) of GO enrichment analysis (P-value < 0.05) in *S. pennellii*. (**C**) The KEGG enrichment scatters plot of cis-regulated target genes of lncRNAs in M82. The size of the circles represents the number of genes, the color of the circle represents the P-value. The abscissa represents the rich factor, the larger the rich factor, the greater the degree of enrichment (RichFactor is the ratio of the number of differentially expressed genes to the number of all genes in this pathway term). (**D**) The KEGG enrichment scatters plot of cis-regulated target genes of lncRNAs in *S. pennellii*.

### 2.5 Construction of lncRNA-mRNA Networks

In order to analyze and compare the functions of lncRNAs and the relationship between lncRNAs and their targeted mRNAs in two tomato cultivars under salt stress, we constructed several putative networks with Cytoscape (Figure 5, S3) As showcased in figures, complex networks were observed. Among them, several genes were found to be involved in oxidation/reduction reaction, phytohormone signaling, and biosynthesis-related in *S. pennellii*, such as ABI2, ACO1, CIPK5/25, CBL4/10, CYP85A1, ABCG25, NCED3/5 and so forth. These 32 mRNAs were predicted to be targeted by 41 lncRNAs (Figure 5). In M82, 13 genes were involved in nitrogen metabolism under salt stress and regulated by 39 lncRNAs (Figure S3). Some transcription factors like MYB were also found to participate in the process of phenotropic biosynthesis. They might activate downstream salt-responsive genes under salt stress conditions. And 25 signal transduction-related genes were also found to be targeted by 78 lncRNAs. Most of them were up-regulated significantly. These relationships among lncRNAs and target genes might play vital roles in sensing and responding to salt stresses.

**Figure 5.**
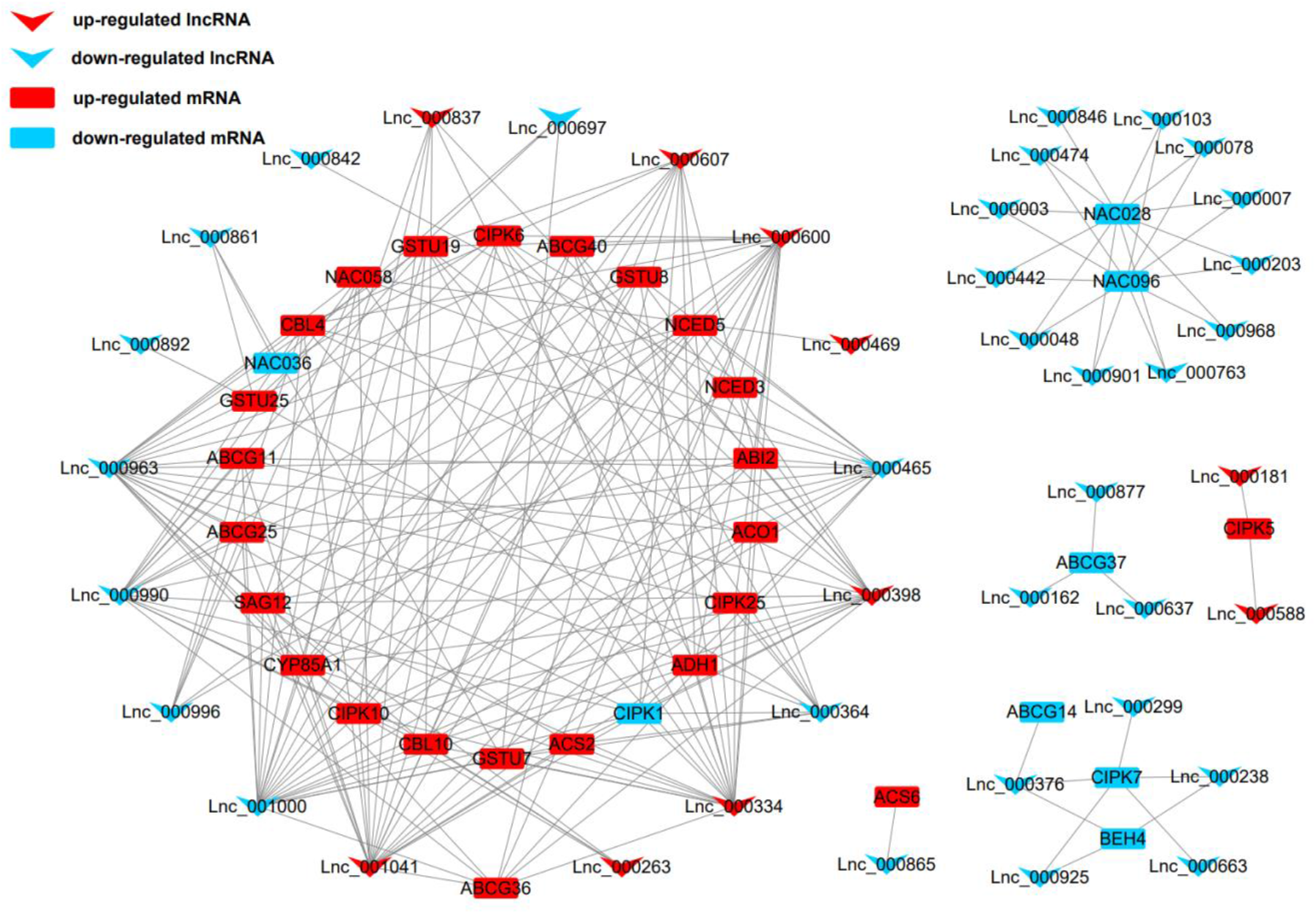
Representatives of the interaction networks among lncRNAs and target protein-coding genes in *S. pennellii*. Red down arrows refer to up-regulated lncRNAs, blue down arrows refer to up-regulated lncRNAs. Red rectangles refer to up-regulated mRNAs, blue rectangles refer to down-regulated mRNAs.

**Figure 6.**
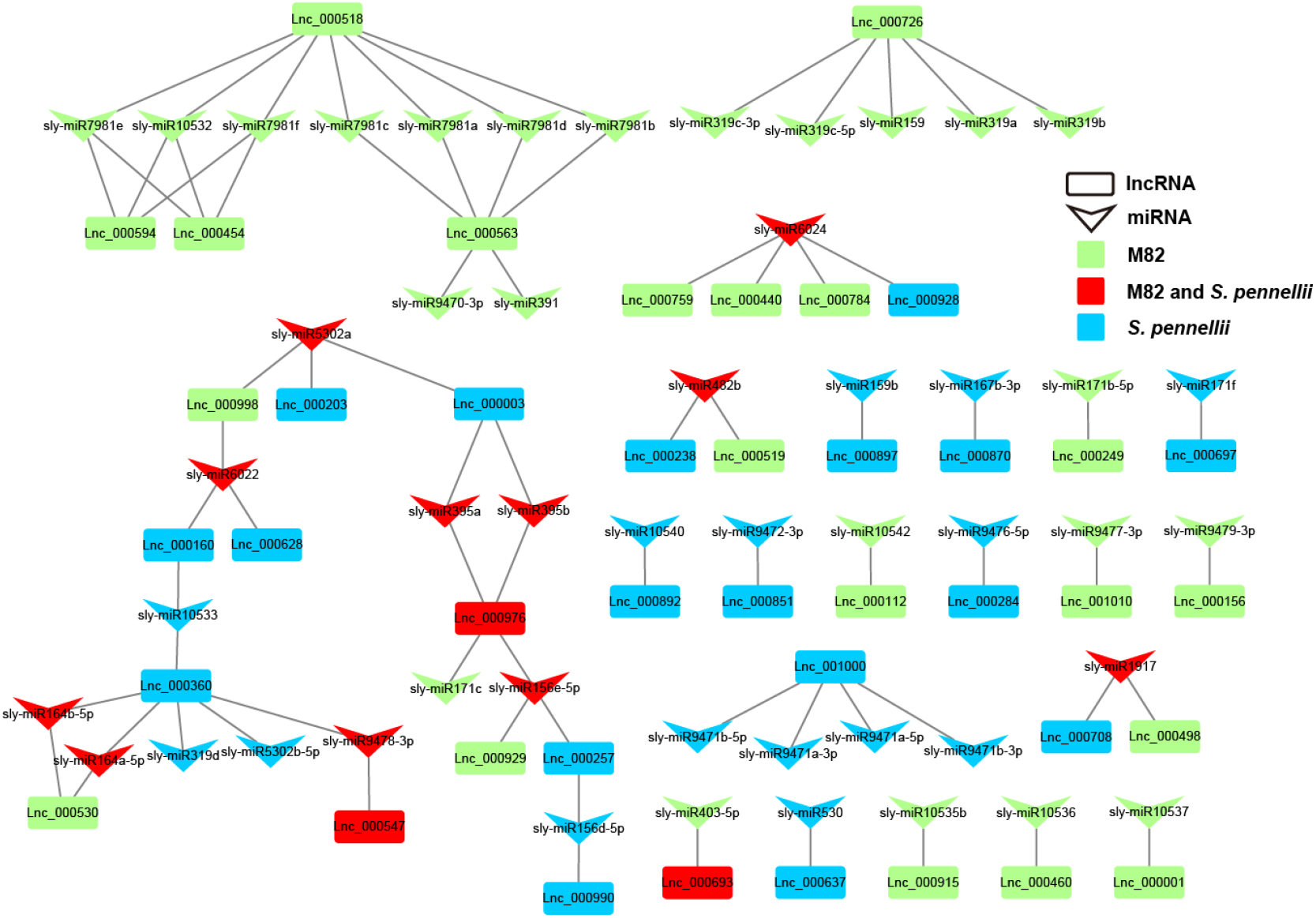
LncRNA–miRNA interaction network. Down arrows refer to miRNAs, rectangles refer to lncRNAs. Green represents the lncRNA that could be targeted by miRNAs only in M82 (or the miRNA that can target DE-lncRNA only in M82). Blue represents the lncRNA that could be targeted by miRNAs only in *S. pennellii* (or the miRNA that can target DE-lncRNAs only in *S. pennellii*). Red represents the lncRNA that could be targeted by miRNAs in both M82 and *S. pennellii* (or the miRNA that can target DE-lncRNAs in both M82 and *S. pennellii*).

### 2.6 The Function of LncRNAs Acting as miRNA Targets in Tomatoes

Previous studies have indicated that the function of lncRNAs could be achieved by targeting miRNAs [20]. Through the psRNATarget server, we screened out the salt-responsive lncRNAs that might be targeted by miRNAs in M82 and *S. pennellii* and established the potential relationships between lncRNAs and miRNAs under salt stress (Supplementary Table S18).

In M82, 23 lncRNAs were predicted to be targets of 34 miRNAs from 24 miRNA families, and 21 lncRNAs were predicted to be targets of 28 miRNAs from 20 miRNA families in *S. pennellii*. There were 3 lncRNAs (Lnc_000547, Lnc_000693, and Lnc_000976) that could be targeted by miRNAs in both two cultivars. Lnc_000547 and Lnc_000693 could be targeted by sly-miR9478-3p and sly-miR403-5p, respectively, while Lnc_000976 could be targeted by 4 miRNAs (sly-miR156e-5p, sly-miR171c, sly-miR395a and sly-miR395b). There were 20 and 18 lncRNAs that could be targeted by the miRNAs in M82 and *S. pennellii*, respectively. In M82, Lnc_000518, Lnc_000563, and Lnc_000726 were the top 3 lncRNAs that connected with miRNAs (7, 6, and 5 miRNAs could be connected respectively). In *S. pennellii*, Lnc_000360, Lnc_000976, and Lnc_001000 were the top 3 lncRNAs that connected with miRNAs (6, 4, and 4 miRNAs could be connected respectively)

Many salt-responsive miRNAs were already discovered in plants, such as miR156 [29], miR159 [30], miR164 [31], miR167 [32, 33], miR171 [34], miR319 [35], miR395 [36] and so forth. In *S. pennelli*, sly-miR156d-5p, sly-miR156e-5p, sly-miR395a, sly-miR395b, sly-miR5302a and sly-miR6022 could target 2 lncRNAs. This illustrated that they could respond to salt stress by the miRNA-lncRNA interactions. Sly-miR156d-5p and sly-miR156e-5p could target Lnc_000990 and Lnc_000976, respectively. Lnc_000257 could be targeted by sly-miR156d-5p and sly-miR156e-5p. Both sly-miR395a and sly-miR395b could target Lnc_000003 and Lnc_000976. Sly-miR5302a could target 2 lncRNAs (Lnc_000003 and Lnc_000203). Sly-miR6022 could target Lnc_000160 and Lnc_000628. While in M82, Lnc_000998 could also be targeted by sly-miR5302a and sly-miR6022. These evidences indicated that lncRNAs could be involved in the gene regulation process through the miRNA– lncRNA interactions under salt stress in tomatoes. The function of these lncRNAs remained to be further determined.

## 3. Discussion

In the past few years, the role of lncRNAs during the normal development process and under abiotic stress in tomatoes has been identified. For instance, the flowering [19], fruit cracking [37], response to drought [23], disease resistance [20], and CRISPR/cas9 technology verified that lncRNA1459 was indeed involved in tomato fruit ripening [38]. With respect to stress-related lncRNAs in other plants, which have been reported according to previous studies. The new lncRNA MuLnc1 in *Morus multicaulis* could regulate mul-miR3954 to produce si161579, which in turn inhibited the expression of MuCML27. The overexpression of MuCML27 could improve the salt stress tolerance of plants [39]. The 1117 salt-responsive lncRNAs in *Gossypium hirsutum* were also reported [26]. In cotton, transgenic plants that increased in seed germination rate, fresh weight, and root length could be obtained by overexpression of lncRNA973, while knockout of lncRNA973 could obtain the plants with more severe wilting and leaf abscission symptoms [40]. There were 126 and 133 DE-lncRNAs were also identified in salt-tolerant and salt-sensitive varieties of *Sweet Sorghum*, respectively. These lncRNAs might play a role as ceRNAs to affect the response to salt stress by regulating the expression of target genes related to ion transport, protein modification, and transcriptional regulation in plants. To the best of our knowledge, this is the first report of the function of lncRNAs in response to salt stress in wild and cultivated tomatoes. The phenotypes of tomatoes under salt stress, such as developmentally arrested seedlings and restricted root growth and development were shown [41]. Therefore, it is important to understand the molecular mechanisms underlying the adaptation of tomatoes to salt stress. As a more salt-tolerant tomato genotype, *S. pennellii* showed relatively more salt responsive compared to cultivated tomato M82. To identify the unique genes that are involved in salt tolerance in wild tomato is essential for cultivating the new salt tolerant tomato cultivars.

In this study, we identified 1044 unique lncRNAs in two cultivars according to strict standards. Some lncRNAs might be excluded during the selection process, but these 1044 lncRNAs could be considered as a group of reliable tomato lncRNAs. Based upon the genome location, these lncRNAs could be divided into four types, including intergenic lncRNAs, antisense lncRNAs, exonic and intronic lncRNAs. We also identified 154 and 137 lncRNAs that were differentially expressed in M82 and *S. pennellii*, respectively. Interestingly, about 86% DE-lncRNAs in M82 were significantly up-regulated, while about 73% DE-lncRNAs in *S. pennellii* were significantly down-regulated. This result revealed a significant difference in the responses of these salt-responsive lncRNAs between cultivated and wild tomatoes under salt stress conditions.

Previous studies showed that lncRNAs had a less conserved sequence [42], it suggested that the DE-lncRNAs in *S. pennellii* might be associated with its high salt tolerance, so the functional elucidation of these DE-lncRNAs mattered deeply. For the most part, lncRNAs can regulate the expression of neighboring genes by cis-regulation and genes located on different chromosomes by trans-regulation [43]. lncRNA can also regulate mRNA expression mediated by miRNAs [44]. Since the salt-tolerance mechanism of *S. pennellii* has not been systematically studied so far. However, lncRNAs have been identified to be an important regulator in response to salt stress in recent years. Therefore, we performed comparative analyses of the salt-responsive lncRNAs target genes between M82 and *S. pennellii*. According to the results we found that some target genes have been confirmed to be related to the salt-tolerance process. In spinach, overexpression of the brassinosteroid (BR) biosynthetic gene CYP85A1 could enhance salt tolerance [45]. Solyc02g089160.3 (CYP85A1) was found to be significantly induced in both two cultivars, indicating that the BR content might be related to salt tolerance in Tomato [46, 47]. As one of the key enzymes from the ethylene (ETH) biosynthetic pathway, ACO1 was significantly upregulated in M82 and could be regulated by five lncRNAs (Lnc_000518, Lnc_000635, Lnc_000693, Lnc_000750, and Lnc_001010). This suggested that these five lncRNAs might play a role in the ethylene synthesis pathway by targeting ACO1 under salt stress. The ACO1 was also involved in the ethylene signal transduction process and could be induced under other stresses [48, 49].

The SALT OVERLY SENSITIVE 3 (SOS3/ CBL4) and SOS3-LIKE CALCIUM BINDING PROTEIN8 (SCABP8/CBL10) help plants cope with Na^+^ toxicity through mediating Ca2^+^ signaling in roots and shoots, respectively [50]. In this study, solyc08g065330.3 (calcineurin B-like protein 10, CBL10), which was trans-regulated by 11 and 13 lncRNAs in M82 and *S. pennellii*, respectively, was significantly up-regulated in both two cultivars. This indicated that CBL10 could be induced ubiquitously in tomatoes under salt stress to help plant scope with Na^+^ toxicity. ADH1 was confirmed to play important roles during stress response in plants [51]. Under salt stress, ADH1 is significantly induced and AtADH1 overexpressing plants showed improved salt stress resistance in Arabidopsis. In this study, ADH1 was significantly up-regulated and could be regulated by 9 and 12 lncRNAs in M82 and *S. pennellii*, respectively.

The plant phytohormones have long been implicated as an important regulator of plant responses to abiotic stress. BR [52, 53], jasmonic acid (JA) [54], gibberellin (GA) [55] and ethylene (ETH) [56] and so forth. We found some genes that might be plant hormone biosynthesis and signal transduction component-encoding genes that were significantly induced in *S. pennellii*. ABA signaling plays an essential role in response to external abiotic stresses. Whereas NCED3, the key gene for ABA synthesis, is considered to be important for the emergence of ABA signals [57]. It can also be regulated by multiple genes, for instance, ATAF1 transcription factor [58], NGTHA1 [59], HDA15 [60]. The expression of NCED3 was significantly induced in *S. pennellii*. As a target gene, Lnc_000842 and Lnc_000996 might be involved in the ABA signaling pathway under salt stress through the transaction on NCED3. In rice, NCED5 was shown to be induced by salt stress. The *nced5* mutant had reduced ABA levels, the ability of the mutant plants to tolerate salt and water stress was also impaired. Furthermore, overexpression of NCED5 could increase ABA levels and enhance tolerability [61]. The expression of NCED5 was significantly induced and predicted to be targeted by 10 lncRNAs. This indicated that lncRNAs might affect ABA levels in *S. pennellii* by regulating the expression levels of NCED3 and NCED5 [62]. AtABCG25 can participate in the intercellular ABA signaling pathway as an ABA transmembrane transporter in *Arabidopsis thaliana* [63], overexpression of AtABCG25 can enhance the ABA accumulation in guard cells and improve plant water use efficiency [64]. Solyc11g018680.1 (AtABCG25) could be significantly induced in *S. pennellii* and targeted by 6 lncRNAs. Besides, AtABCG25 can also transport ABA together with AtABCG40 to achieve stomatal regulation [65, 66]. Solyc09g091670.3 (AtABCG40) was predicted to be targeted by 6 and 7 lncRNAs in M82 and *S. pennellii*, respectively. It was significantly up-regulated in two cultivars. In contrast to M82, lncRNAs might enhance ABA signaling transduction by targeting AtABCG25 and AtABCG40 simultaneously to adapt to salt stress via more rapid stomatal regulation. As a synergid-expressed kinase, FERONIA plays a critical role in hormone signaling and stress tolerance in different plants [67]. While the FERONIA activity could be regulated by ABI2 [67]. In *S. pennellii*, ABI2 was significantly induced. 10 lncRNAs might mediate FERONIA signaling pathway by cis-regulating ABI2 expression. Ethylene is an important plant hormone involved in plant growth, development and response to environmental stresses [68]. 1-aminocyclopropane-1-carboxylic acid synthase (ACS) is a key rate-limiting enzyme responsible for ethylene biosynthesis in plants [69]. And the ethylene synthesis pathway is tightly regulated by exogenous and endogenous signals at the transcriptional and post-transcriptional levels. MAPK phosphorylation-induced stabilization of ACS6 protein is mediated by the non-catalytic C-terminal domain, which also contains the cis-determinant for rapid degradation by the 26S proteasome pathway. In *Arabidopsis*, as the MAPK substrates, ACS2 and ACS6 could be phosphorylated by MPK6 and results in ACS accumulation and induction of ethylene [70, 71]. ACS6 was found to be induced by Lnc_000865 only in *S. pennellii*, while ACS2 could be induced in both two cultivars. It showed that ACS2 might be necessary under salt stress to initiate ethylene biosynthesis in tomatoes, while the up-regulated ACS6 might enhance biosynthesis of ethylene to improve salt tolerance of *S. pennellii*. Previous study showed that DWF4, as a BR biosynthesis gene, could be directly inhibited by BZR1 and that its expression level increased at an early stage of stress [72, 73]. A homolog of BZR1 called BEH4 was found to be targeted by 3 lncRNA and inhibited in *S. pennellii*. As the target gene of BZR1, DWF4 was significantly up-regulated. It showed that lncRNAs might mediate the BR biosynthesis process to enhance the salt tolerance in *S. pennellii*. Previous studies have shown that high[Ca2^+^] cyt can be sensed by (CBLs). However, CBL4 and CBL10, a SOS3 like calcium-binding protein, can help plants cope with Na^+^ toxicity by mediating Ca2^+^ signals in roots and shoots, respectively. Act as the main Ca2^+^ sensor, CBLs can bind Ca2^+^ and regulate the activity of many proteins [50, 74, 75]. Solyc12g055920.2 (CBL4) could be induced in *S. pennellii* and no significant changes were noted in M82. In *Arabidopsis*, CBL4 can interact with CIPK6 to regulate the activity of the K^+^ channel protein (AKT2) [76]. CBL4 was also shown to interact with CIPK5 in bermudagrass, and overexpression of either CBL4 or CIPK5 could increase salt tolerance [77]. Solyc08g067310.1 (CIPK5) could be targeted by 2 lncRNAs (Lnc_000181 and Lnc_000588) induced in *S. pennellii*, while Solyc12g010130.1 (CIPK6) could be targeted by 9 lncRNAs and induced only in *S. pennellii*. It showed that lncRNAs in *S. pennellii* might help plants cope with Na^+^ toxicity by regulating Ca2^+^ signaling in the root rapidly. Tau class glutathione transferases (GSTU) genes could protect plants from oxidative injury, overexpression of SbGST showed higher germination rates [78, 79]. In this study, Solyc06g069045.1 (GSTU7) and Solyc09g011510.2 (GSTU8) were both induced in *S. pennellii*. Previous studies showed that NAC proteins was involved in plant salt stress response [80, 81]. In Reaumuria trigyna, NAC100 could interact with RtRbohE/SAG12 to accelerate salt-induced programmed cell death. NAC100 promoted reactive oxygen species, Ca2^+^, and Na^+^ accumulation and increased the Na^+^/K^+^ ratio [82]. Interestingly, both NAC100 and SAG12 were differentially up-regulated in *S. pennellii* and 10 lncRNAs could target these two genes. Furthermore, Lnc_000364 and Lnc_000263 could target NAC100 and SAG12, respectively. It illustrated that *S. pennellii* might adapt to the salt stress environment by regulating the programmed death process of cells via lncRNAs.

Moreover, one of the molecular mechanisms by which lncRNAs regulate the expression of genes is to interact with miRNAs as ceRNAs [23]. By using psRNAtarget, 41 lncRNAs were predicted as potential targets of 49 miRNAs. In particular, miR156 was confirmed to be associated with salt stress, overexpression of miR156a could reduce salt tolerance of apple [83]. In maize, zma-miR159 was up-regulated under salt stress, while mir159 could affect the ABA signaling regulatory process through interacting with its target gene MYB. Whereas zma-miR164 could respond to the salt stress by targeting the NAC transcription factors [84]. In *Populus euphratica*, peu-miR164 was predicted to target PeNAC070. Overexpression of NAC070 could reduce salt tolerance in *Arabidopsis* [85]. It indicated that miR164 is a positive regulator of the salt tolerance pathway in Populus euphratica. MiR319 plays important role in response to abiotic stress in some C3 plants, like *Arabidopsis* and rice, for example. In Panicum virgatum L, where researchers found that miR319 could modulate the salt response of switchgrass by fine-tuning ET synthesis [35]. Different lncRNAs were specifically targeted by these salt-related miRNAs in both two cultivars. In *S. pennellii*, Lnc_000257 and Lnc_000990 could be targeted by sly-mir156d-5p, Lnc_000257 and Lnc_000976 could be targeted by sly-miR156e-5p. Lnc_000360 could be targeted by 3 miRNAs (sly-miR164a-5p, sly-miR164b-5p and sly-miR319d). While in M82, Lnc_000929 和 Lnc_000976 could be targeted by sly-miR156e-5p, sly-miR164a-5p and sly-miR164b-5p could target Lnc_000530. These results suggested that these lncRNAs might respond to the salt stress process by interacting with miRNAs and had significant differences between the two tomato cultivars.

In summary, we found that some pathways such as phytohormone metabolism, photosynthesis, protein/amino acid metabolism were closely related to salt stress by analyzing the function of lncRNA target genes in M82 and *S. pennellii*. Phytohormones regulate many growths, developmental processes, and responses to various stresses. Following the target genes prediction, we obtained some lncRNAs that were involved in phytohormone biosynthesis and transport pathways. Under salt stress, the expression levels of some genes involved in hormone synthesis including ABA and ethylene were significantly up-regulated in *S. pennellii*, and the mean expression levels were also higher than that in M82, which might account for the differences in salt tolerance of cultivated and wild genotypes. Increased cell death-related genes expression in *S. pennellii* might result in having the chance to receive more death signals to induce apoptosis and contributed to the tolerance to salt stress. In addition, we also analyzed the relationships between salt-responsive lncRNAs and miRNAs to elucidate the roles and distinction of salt-responsive lncRNAs in wild and cultivated tomatoes from several different standpoints. We also constructed several putative salt tolerance-related networks that were associated with the high salt tolerance of *S. pennellii*. In particular, we will expect to analyze the functions of these salt-responsive lncRNAs by utilizing more scientifically rigorous methods.

## 4. Materials and Method

### 4.1 Plant Materials and Stress Treatment

Seeds of cultivated tomato M82 (salt-sensitive) and wild tomato *S. pennellii* (salt-tolerant) [86] were sown in pots containing 3:1 mixtures of vermiculite: perlite (V/V) in the growth chamber under a 16 h/8 h (day/night), and a light intensity of 100 μmol m ^−2^ sec-1, 25 °C, 20-30% relative humidity. Plants were cultivated for 6 weeks (well-watered and with Hoagland solution supplying at an interval of 2 weeks), and then the seedlings were exposed to 200 mM NaCl (salinity). Plants grown in the same environment without the additional stress component were used as controls. Roots of tomatoes were collected 0 h and 12 h following exposure to stress.

### 4.2 RNA Extraction, Construction of cDNA Libraries and High-Throughput Sequencing

Total roots RNA of tomatoes was isolated using RNAprep pure Plant Kit (Tiangen Biotech, Beijing, China) according to the manufacturer’s protocol. RNA quality and quantity were checked with NanoDrop 2000 Spectrophotometer (Thermo Fisher Scientific, Wilmington, USA) and an Agilent Bioanalyzer 2100 System (Agilent Technologies, CA, USA), respectively. RNA samples were pooled with equal amounts of RNA from three independent individuals. The samples were sent to Biomarker Technologies Co. Ltd. (Beijing, China) for lncRNAs libraries preparation and Illumina sequencing. Poly(A) RNA enrichment and strand-specific RNA-seq library were prepared to use the NEBNext^®^ UltraTM RNA Library Prep Kit for Illumina (NEB, USA) according to the low sample protocol guidelines. Libraries were controlled for quality using the NanoDrop 2000 Spectrophotometer. The resulting libraries were sequenced on an Illumina sequencing platform with paired-end reads of 150 bp.

### 4.3 RNA-Seq Reads Mapping and Transcriptome Assembling

FastQC was utilized to check the quality of RNA-seq data. And then the adapters of raw reads that capture at least one of the following two characteristics: more than 20% of bases with a Q-value≤20 or an ambiguous sequence content (“N”) exceeding 5% were removed. After aligning the clean reads to the reference genome by using HISAT2 [87] with default settings, the reads were assembled by using Cufflinks [88].

### 4.4 Identification of Salt-Responsive LncRNAs

To identify putative lncRNA transcripts, all the mRNA transcripts were filtered out firstly. Then the transcripts with a length less than 200 nt or only have one exon were also filtered out. Then, the protein-coding ability of the remaining transcripts was predicted by using the CPC [89], PLEK [90], and CNCI [91] software. The Swiss-Prot database was also used to filter the transcripts with any known domains [92]. The transcripts without protein-coding ability were subsequently employed in the remainder of the study. LincRNAs, antisense lncRNAs, intronic lncRNAs and exonic lncRNAs were classified by the cuffcompare software.

### 4.5 Analysis of Differentially Expressed LncRNAs

The expression levels of lncRNAs were normalized by Transcripts Per Million (TPM). Then the R package DESeq2 was utilized to perform the differentially expressed analysis [93]. The fold changes of lncRNAs were calculated via log_2_(TPM). The lncRNAs exhibiting a |fold change|≥2 and P-value < 0.05 were considered the DE-lncRNAs.

### 4.6 Prediction of DE-lncRNAs Target Genes and Analysis of LncRNAs Function

The functional annotation of DE-lncRNAs was carried out by co-location and co-expression analysis. The co-locating and co-expressing lncRNA-mRNA pairs were identified by custom Perl scripts. The coding genes 1000k upstream and downstream of lncRNAs were considered to be co-located.

### 4.7 GO and KEGG Enrichment Analysis

The functions of the target genes of DE-lncRNAs were annotated by GO and KEGG enrichment analysis. The GO enrichment analysis was performed on the agrigo website (http://systemsbiology.cau.edu.cn/agriGOv2/). The KEGG analysis was performed on the KOBAS website (http://kobas.cbi.pku.edu.cn/kobas3).

### 4.8 Interaction Analyses of lncRNA-miRNA and miRNA-mRNA Pairs

A total of 147 known miRNA sequences of tomato were downloaded from the miRBase database (https://www.mirbase.org) [94] and were utilized for analyzing the interaction relationship between lncRNAs and miRNAs by using the website tool psRNATarget (http://plantgrn.noble.org/psRNATarget/) with the default parameters [95]. The interaction relationships between miRNAs and mRNAs were also implemented by the methods above for description and parameters. Finally, according to the ceRNA regulatory mechanism and the relationships between lncRNA-miRNA and miRNA-mRNA pairs, the ceRNA regulatory networks were constructed and visualized by Cytoscape software (v3.7.2) [96].

## Supplementary Materials

Table S1: RNA-seq data quality assessment; Table S2: expression levels of all identified lncRNAs; Table S3: Differentially expressed lncRNAs in M82; Table S4: Differentially expressed lncRNAs in *S. pennellii*; Table S5: cis-regulated lncRNA-mRNA pairs in M82; Table S6: cis-regulated lncRNA-mRNA pairs in *S. pennellii*; Table S7: GO enrichment results of cis-regulated target genes pairs in M82; Table S8: GO enrichment results of cis-regulated target genes pairs in *S. pennellii*; Table S9: KEGG enrichment results of cis-regulated target genes pairs in M82; Table S10: KEGG enrichment results of cis-regulated target genes pairs in *S. pennellii*; Table S11: trans-regulated lncRNA-mRNA pairs in M82; Table S12: trans-regulated lncRNA-mRNA pairs in *S. pennellii*; Table S13: GO enrichment results of trans-regulated target genes pairs in M82; Table S14: GO enrichment results of trans-regulated target genes pairs in *S. pennellii*; Table S15: GO enrichment results of trans-regulated target genes pairs in both M82 and *S. pennellii*; Table S16: KEGG enrichment results of trans-regulated target genes pairs in M82; Table S17: KEGG enrichment results of trans-regulated target genes pairs in *S. pennellii*; Table S18: KEGG enrichment results of trans-regulated target genes pairs in both M82 and *S. pennellii*; Figure S1: GO enrichment results of trans-regulated target genes pairs in both M82 and *S. pennellii*; Figure S2: KEGG enrichment results of trans-regulated target genes pairs in both M82 and *S. pennellii*; Figure S3: The relationships between DE-lncRNAs and DE-mRNAs in M82.

## Author Contributions

Conceptualization, J.G. and H.W.; Methodology, N.L. and Z.W.; Software, T.Y. and S.H.; Validation, B.W. and J.W.; Formal analysis, N.L. and Z.W.; Investigation, N.L. and Z.W.; Resources, Q.Y. and J.G.; Data curation, N.L. and Z.W.; Writing - original draft, N.L. and Z.W.; Writing - review & editing, N.L. and Z.W.; Visualization, N.L. and R.X.; Supervision, J.G. and H.W.; Project administration, J.G. and H.W.; Funding acquisition, J.G. and H.W.. All authors have read and agreed to the published version of the manuscript.. All authors have read and agreed to the published version of the manuscript.

## Data Availability Statement

The raw sequence data reported in this paper have been deposited in the Genome Sequence Archive (Genomics, Proteomics & Bioinformatics 2021) in National Genomics Data Center (Nucleic Acids Res 2021), China National Center for Bioinformation / Beijing Institute of Genomics, Chinese Academy of Sciences, under accession number CRA004289 that is publicly accessible at https://ngdc.cncb.ac.cn/gsa.

## Acknowledgments

The authors thank the reviewers and all of the editors for their helpful comments and suggestions on this paper.

