## Supplementary figures and images for "Identification and Characterization of Long Non-Coding RNA in Tomato Roots under Salt Stress"

### supplemental Figure S1.pdf

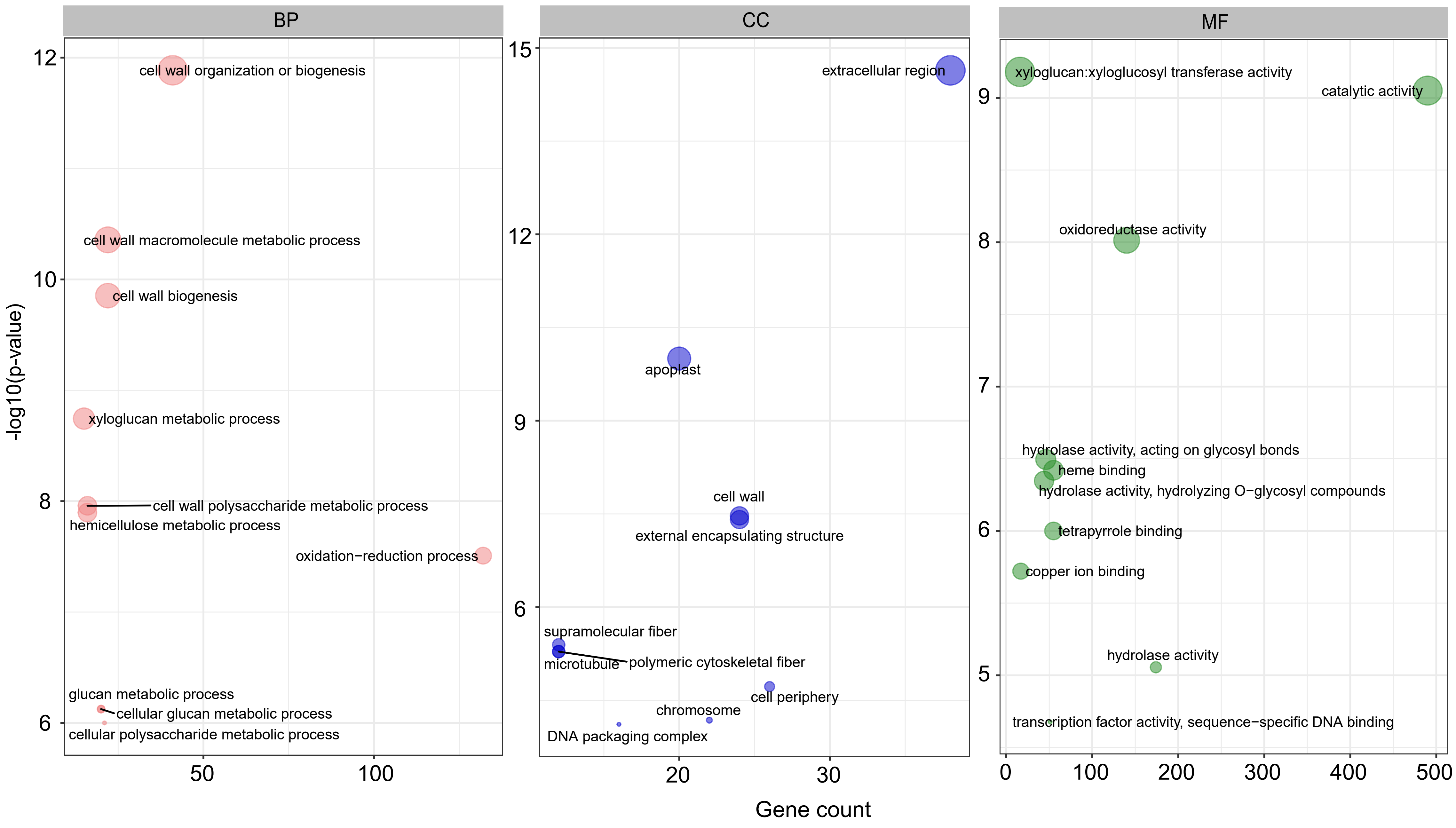

### supplemental Figure S2.pdf

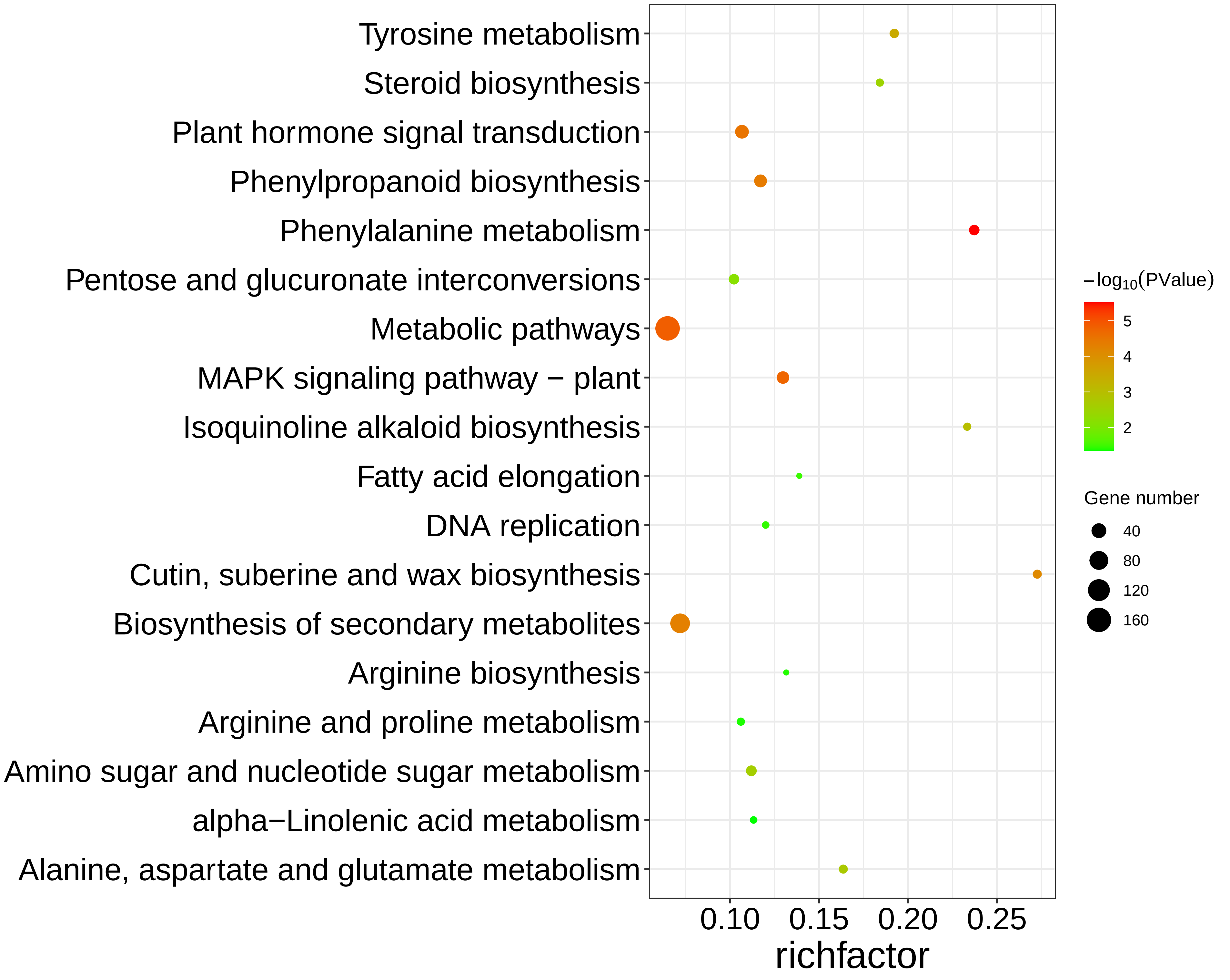

### supplemental Figure S3.pdf

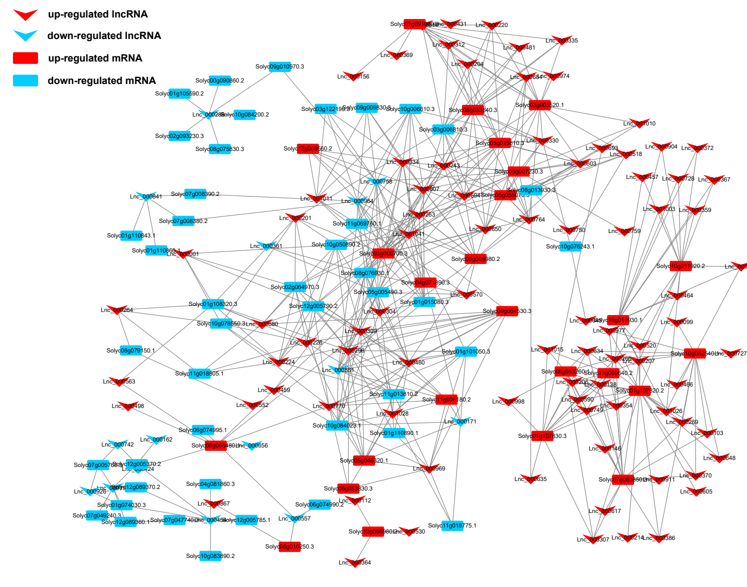
